# Evolving a new efficient mode of fructose utilization for improved bioproduction in *Corynebacterium glutamicum*

**DOI:** 10.1101/2021.02.18.431779

**Authors:** Irene Krahn, Daniel Bonder, Lucia Torregrosa, Dominik Stoppel, Jens P. Krause, Natalie Rosenfeldt, Tobias M. Meiswinkel, Gerd M. Seibold, Volker F. Wendisch, Steffen N. Lindner

## Abstract

Fructose utilization in *Corynebacterium glutamicum* starts with its uptake and concomitant phosphorylation via the phosphotransferase system (PTS) to yield intracellular fructose 1-phosphate, which enters glycolysis upon ATP dependent phosphorylation to fructose 1,6-bisphosphate by 1-phosphofructokinase. This is known to result in a significantly reduced oxidative pentose phosphate pathway (oxPPP) flux on fructose (~10 %) compared to glucose (~60 %). Consequently, the biosynthesis of NADPH demanding products, e.g. L-lysine, by *C. glutamicum* is largely decreased, when fructose is the only carbon source. Previous works reported that fructose is partially utilized via the glucose specific PTS presumably generating fructose 6-phosphate. This closer proximity to the entry point of the oxPPP might increase oxPPP flux and consequently NADPH availability. Here, we generated deletion strains either lacking in the fructose-specific PTS or 1-phosphofructokinase activity. We used these strains in short-term evolution experiments on fructose minimal medium and isolated mutant strains, which regained the ability of fast growth on fructose as a sole carbon source. In these fructose mutants, the deletion of the glucose specific PTS, as well as the 6-phosphofructokinase gene, abolished growth, unequivocally showing fructose phosphorylation via glucose specific PTS to fructose 6-phosphate. Gene sequencing revealed three independent amino acid substitutions in PtsG (M260V, M260T, P318S). These three PtsG variants mediated faster fructose uptake and utilization compared to native PtsG. In-depth analysis of the effects of fructose utilization via these PtsG variants revealed significantly increased biomass formation, reduced side-product accumulation, and increased L-lysine production by 50 %.

## Introduction

Canonical metabolic routes evolved for superior performance in the natural habitat, but often they do not represent the ideal choice from a biotechnological perspective (Erb et al., 2017). If more suitable alternative pathways are known, rational approaches of metabolic engineering can redirect metabolic pathways into more advantageous directions. In the absence of a known and better suited natural alternative, adapted laboratory evolution (ALE) may select for efficient pathway variants. *Corynebacterium glutamicum* is employed in the million-ton scale bioproduction of amino acids, with the lion’s share split between L-glutamate and L-lysine (Wendisch, 2020). Beyond amino acids, amines, organic acids and alcohols are produced with this bacterium (Becker et al., 2018;Mindt et al., 2020).

NADPH is an important cofactor for anabolic reactions, and hence a limiting factor in the production of metabolites with a particularly high demand for NADPH, e.g. L-lysine, which requires 4 molecules NADPH per molecule L-lysine produced (Marx et al., 1996). To provide NADPH, *C. glutamicum* possesses several dehydrogenases, which use NADP cofactor. These are the glucose 6-phosphate dehydrogenase (Zwf), and the 6-phosphogluconate dehydrogenase (Gnd) of the oxidative part of the pentose phosphate pathway (oxPPP), the isocitrate dehydrogenase (Icd) in the TCA cycle, and the malic enzyme (MalE), and their overexpression improved production L-lysine (Georgi et al., 2005;Becker et al., 2007). NADPH provision was optimized by heterologous expression of genes encoding the membrane-bound transhydrogenase from *E. coli* (Kabus et al., 2007). Although *C. glutamicum* lacks transhydrogenase, it is known to run an ATP consuming transhydrogenase-like cycle between the anaplerotic reactions, malate dehydrogenase and malic enzyme, transferring electrons from NADH to NADP (Blombach et al., 2011). However, the predominant way of NADPH generation is via the oxPPP.

Metabolic engineering was used to broaden the substrate spectrum of *C. glutamicum* towards second generation feedstocks such as non-food wastes from biodiesel (glycerol) (Rittmann et al., 2008) or hemicellulose biomasses (xylose, arabinose) (Zhao et al., 2018). Yet, the sugars glucose, derived from starch hydrolysates as well as sucrose and fructose derived from molasses are still the preferred carbon sources for amino acid production. The decrease in L-lysine yields when fructose is used instead of glucose is drastic (Kiefer et al., 2004;Georgi et al., 2005). Although the entry points of glucose and fructose are only two reactions apart, the fluxes through the oxPPP, and hence the prevalent NADPH generating reactions, are significantly different. On fructose a very low flux is described (10 %), whereas glucose leads to a high flux (60 %) (Kiefer et al., 2004). As result a low L-lysine product yield is reached on fructose compared to glucose (Georgi et al., 2005). Moreover, the reported high oxPPP fluxes on glucose may still be limiting for overproduction of high NADPH consuming products, such as amino acids (Murai et al., 2020).

The only known way for *C. glutamicum* to phosphorylate fructose and sucrose, and hence turn them into central metabolism intermediates, is via the phosphoenolpyruvate-dependent phosphotransferase system (PTS), for glucose on the other hand, an ATP pathway is present (Moon et al., 2007;Lindner et al., 2011) (Ikeda, 2012). Glucose is phosphorylated to glucose 6-P by glucose specific PTS compound (PtsG). Sucrose is phosphorylated to sucrose 6-P via its PTS (PtsS) and subsequently cleaved to glucose 6-P and fructose. Fructose, regardless if added to the medium as a carbon source or originating from sucrose catabolism, is taken up and phosphorylated to fructose 1-P by a fructose specific PTS (PtsF) (Dominguez and Lindley, 1996;Parche et al., 2001). After a second ATP dependent phosphorylation of fructose 1-P catalyzed by 1-phosphofructokinase (FruK), it enters glycolysis at the level of fructose 1,6-BP. Additionally a minor fraction of fructose (< 10 %) is taken up and phosphorylated by PtsG to generate fructose 6-phosphate (Kiefer et al., 2004) (Figure 1). Most strikingly, overexpression of fructose 1,6-bisphosphatase increased L-lysine production when fructose was used as the carbon source (Georgi et al., 2005), pointing to an advantage of fructose 6-P over fructose-1,6-BP for increasing oxPPP-flux and, consequently, increasing NADPH regeneration and productivity. Thus, shifting the carbon flux slightly closer to the entry-point of the oxPPP allows a higher flux through the oxPPP and a higher NADPH regeneration rate. Similarly, L-lysine production from molasses was optimized by overexpression of fructose 1,6-bisphosphatase and fructokinase (Xu et al., 2013).

**Figure 1.**
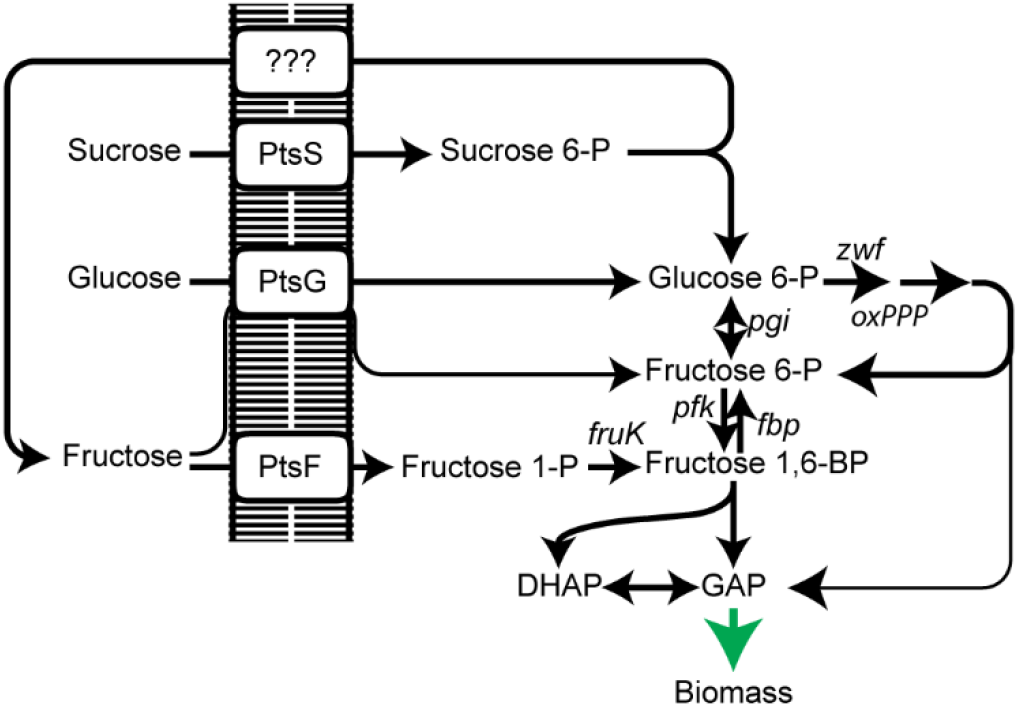
Scheme of PTS-dependent uptake and utilization of sugars sucrose, glucose and fructose in *C. glutamicum*. PtsS, sucrose PTS; PtsG, glucose PTS; PtsF, fructose PTS; oxPPP, oxidative pentose phosphate pathway; *pgi*, phosphoglucose isomerase; *pfk*, 6-phosphofructokinase; *fbp*, fructose 1,6-bisphosphatase; *fruK*, 1-phosphofruktokinase; DHAP, dihydroxyacetone phosphate; GAP, glyceraldehyde 3-phosphate.

Here we aimed on increasing the efficiency of the PtsG catalyzed conversion of fructose to fructose 6-P. We generated strains unable to utilize fructose via its usual route and selected fast growing strains after short-term evolution in fructose minimal medium. Isolated PtsG variants were identified and reverse engineering complemented fructose utilization in the deletion strains. Deletion of either 6-phosphofructokinase in the mutants and overexpression of the PtsG variants in a fructose 1,6-bisphosphatase deletion strain confirmed fructose phosphorylation to fructose 6-P by the PtsG variants. ^13^C-labeling experiments revealed that a higher oxPPP flux is present in the reverse engineered strains. Finally, the new way of fructose utilization was tested on L-lysine production, showing an increase in L-lysine yield from fructose.

## Materials and Methods

### Strains and plasmids used

*C. glutamicum* strains and plasmids used are listed in Table 1 and Table 2, respectively. For plasmid construction the primers listed in Supplementary Table S1 were used. For cloning genes were amplified from genomic DNA and cloned by the indicated restriction sides (Supplementary Table S1) into similarly restricted pVWEx1. Deletion plasmids were constructed by cloning PCR-fused products of primer pairs A+B and C+D and cloned blunt-ended into *Sma*I digested pK19mobsacB.

**Table 1.**
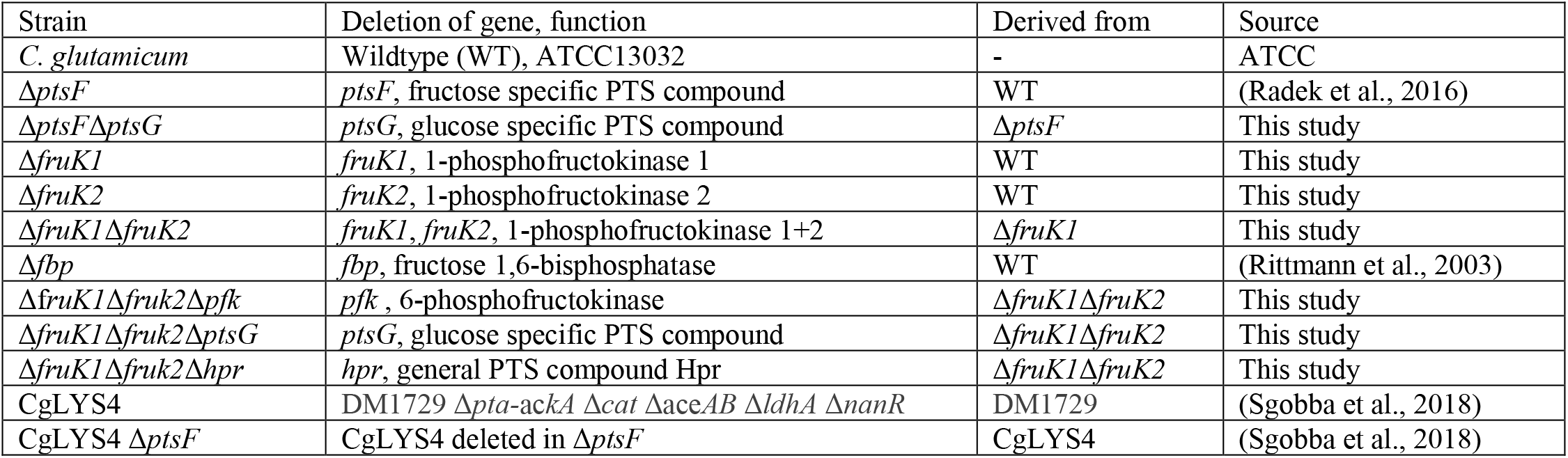
*C. glutamicum* strains used in this study.

**Table 2.** Plasmids used in this study,

| name | Properties / use | source |
| --- | --- | --- |
| pVWEx1 | Km<sup>R</sup>, pHM1519, *P<sub>tac</sub>*, *lacI<sup>q</sup>*, for IPTG inducible expression | (Peters-Wendisch et al., 2001) |
| pVWEx1-*ptsG* | pVWEx1, carrying WT *ptsG* | This study |
| pVWEx1-*ptsG*<sup>M260V</sup> | pVWEx1, carrying *ptsG* mutated in M260V | This study |
| pVWEx1-*ptsG*<sup>M260T</sup> | pVWEx1, carrying *ptsG* mutated in M260T | This study |
| pVWEx1-*ptsG*<sup>P318S</sup> | pVWEx1, carrying *ptsG* mutated in P318S | This study |
| pVWEx1-*ptsF* | pVWEx1, carrying WT *ptsF* | This study |
| pVWEx1-*lysC*<sup>fbr</sup> | pVWEx1, carrying feedback resistant version of *lysC* (T311I) | This study |
| pK19*mobsacB* | Km<sup>R</sup>, RP4; *mob*; oriV<sub>Ec</sub>; *sacB*; *lacZα*; for allelic exchange | (Schafer et al., 1994) |
| pK19*mobsacBΔfruK1* | pK19*mobsacB* based for *fruK1* deletion | This study |
| pK19*mobsacBΔfruK2* | pK19*mobsacB* based for *fruK2* deletion | This study |
| pK19*mobsacBΔpfk* | pK19*mobsacB* based for *pfk* deletion | This study |
| pK19*mobsacBΔptsG* | pK19*mobsacB* based for *ptsG* deletion | This study |
| pK19*mobsacBΔptsF* | pK19*mobsacB* based for *ptsF* deletion | (Radek et al., 2016) |
| pK19*mobsacBΔhpr* | pK19*mobsacB* based for *hpr* deletion | (Lindner et al., 2011) |

### Culture conditions & growth experiments

*C. glutamicum* strains were cultivated in LB (0.5 % NaCl, 1 % tryptone, 0.5 % yeast extract) or CgXII minimal medium (20 g/L (NH_4_)_2_ SO_4_, 5 g/L urea, 1 g/L KH_2_PO_4_, 1 g/L K_2_HPO_4_, 42 g/L MOPS, 10 mg/L CaCl_2_, 250 mg/L MgSO_4_ × 7 H_2_O, 0.01 mg /L FeSO_4_ × 7 H_2_O, 0.01 mg/L MnSO_4_ × 7 H_2_O, 0.001 mg/L ZnSO_4_ × 7 H_2_O, 0.0002 mg/L CuSO_4_, 0.00002 mg/L NiCl_2_ × 6 H_2_0, pH 7) (Eggeling and Bott, 2005). For growth experiments the strains grew in 50 ml LB cultures overnight, harvested by centrifugation (3.220 \**g*), washed twice in CgXII w/o carbon source and inoculated to an optical density of 1 in 50 ml CgXII containing the indicated carbon sources. For plasmid construction, *E. coli* DH5α was used and cultured in LB medium (1 % tryptone, 0.5 % yeast extract, 1 % NaCl). Precultures for growth experiments with *C. glutamicum* and all *E. coli* cultures were carried out in LB. For selection on pVWEx1 and derivatives, 50 and 25 mg/mL kanamycin was added to *E. coli* and *C. glutamicum* cultures, respectively. CgXII minimal medium (Eggeling and Bott, 2005) was used for growth, sugar uptake, and L-lysine production experiments. Cells were harvested in the exponential growth phase by centrifugation (RT, 3,220 *g for 10 min) and washed twice in CgXII medium without carbon source. Gene-expression was induced by addition of up to 1 mM isopropyl β-D-1-thiogalactopyranoside (IPTG). Ideal concentration of IPTG for *ptsF*/*G* expression was determined to be at 30 µM IPTG. Cultivations were carried out in 50 mL solutions in 500 mL baffled shaking flasks at 120 rpm and 30°C.

### Analysis of sugars and organic acids concentration, and amino acid production

Lysine production: To verify L-lysine production, strains were inoculated to OD_600_ of 1 in CgXII media supplemented with 4 % fructose, 30 µM IPTG and, if carrying a pVWEx1 variant, 50 µg/mL kanamycin in 500ml baffled shake flasks. Supernatants were collected at 4, 8, 12, 24, 48, and 72h after inoculation. L-lysine concentrations were determined in up to 1:5000 serial dilution of supernatants using an ICS-6000 HPIC Ion Chromatography equipped with an AminoPac PA10 IC column, ICS-6000 CD Conductivity Detector, and ADRS 600 Anion Dynamically Regenerated Suppressor (Dionex, CA, USA). The column was set with a 10 mM - 250 mM NaOH gradient at a 0.25ml/min flow rate. Sugars and organic acid concentrations were quantified via HPLC as described previously (Rittmann et al., 2008).

### ^13^C isotopic labelling of proteinogenic amino acids

^13^C-isotope tracing was performed to indirectly analyze carbon flux. Cells were cultured in 4 mL CgXII medium containing ^13^C-1-glucose or ^13^C-1-fructose (Sigma-Aldrich, Taufkirchen, Germany) as sole carbon sources. Cultures were inoculated from CgXII + 20 mM pyruvate overnight cultures to an OD_600_ 0.01 and grown at 30°C until early stationary phase. Before inoculation cells were washed twice (RT, 6000\**g*, 3 min) in carbon source free CgXII medium. 10^9^ cells (~1 mL of OD_600_ = 1) were pelleted, washed with ddH_2_O and hydrolyzed in 1 mL 6N hydrochloric acid at 95°C for 24 h. Subsequently to hydrolysis HCl was evaporated by heating at 95°C under an air-stream. Hydrolyzed biomass was resuspended in 1 mL ddH_2_O. Amino acids masses were analyzed after separation by ultra-performance liquid chromatography (Acquity, Waters, Milford, MA, USA) using a C18-reversed-phase column (Waters, Eschborn, Germany) as previously described (Giavalisco et al., 2011). Mass spectra were acquired by an Exactive mass spectrometer (Thermo Scientific, Dreieich, Germany). Data was analyzed using Xcalibur (Thermo Scientific, Dreieich, Germany). Amino-acid standards (Merck, Darmstadt, Germany) were used to determine specific retention times.

### Sugar uptake measurements

For ^14^C-labeled fructose uptake studies, strains were grown to early exponential growth phase with 50 mM fructose as sole carbon source and 30 µM IPTG, if appropriate. Cells were harvested by centrifugation, washed two times in ice-cold CgXII medium (without carbon source), resuspended to an optical density OD_600_ of 2 in CgXII medium and stored on ice until measurement. Prior to the transport assay, cells were incubated for 3 min at 30°C. The assay was started by addition of 1 µM to 1 mM ^14^C-labeled fructose (specific activity of 45 mCi mmol^−1^; Hartmann Analytik, Braunschweig, Germany). At given time intervals (15, 30, 45, 60, and 120 s), 200 µl samples were filtered through glass fiber filters (type F; Millipore, Eschborn, Germany) and washed twice with 2.5 ml of 100 mM LiCl. The radioactivity of the filter samples was determined using scintillation fluid (Rotiszinth; Roth, Germany) and a scintillation counter (LS 6500; Beckmann, Krefeld, Germany).

## Results

### Construction and characterization of strains lacking fructose specific PTS and 1-phosphofructokinase genes

Previous studies suggested a PtsG mediated fructose utilization to fructose 6-P in *C. glutamicum* (Kiefer et al., 2004). We assumed that the direct generation of fructose 6-P instead of fructose 1,6-BP from fructose increases oxidative PPP flux leading to higher NADPH-availability, which is advantageous for high NADPH-demanding bioproductions, such as L-lysine. In fact, overexpression of fructose 1,6-bisphosphatase increased L-lysine production from fructose (Georgi et al., 2005). The aim of this study was to explore the promiscuous reaction of PtsG and evolve it for increased activity.

To be able to select for this route of fructose utilization and to evolve it, we generated two strains, which are deficient in the canonical route of fructose utilization. The first lacks the fructose specific PTS compound (∆*ptsF*), and the second lacks 1-phosphofructo kinase activity (∆*fruK1* ∆*fruK2*). As expected, growth of these strains on fructose was strongly affected. The ∆*ptsF* strain grew with a very low growth rate and the ∆*fruK1* ∆*fruK2* strain did not grow at all within 24 h (Figure 2). Thus, both strains were considered suitable for performing shake-flask short-term evolution experiments. In particular, the slow growth of ∆*ptsF* indicates the presence of an alternative way for fructose utilization in our background-strain, suggesting a good starting-point for optimization of the reaction through evolution. In contrast to the described results, a deletion strain lacking the general PTS compound HPR is unable to grow on fructose, also after prolonged incubation, pointing to the contribution of the earlier reported fructose uptake via PtsG (Kiefer et al., 2004;Moon et al., 2007) (data not shown).

**Figure 2.**
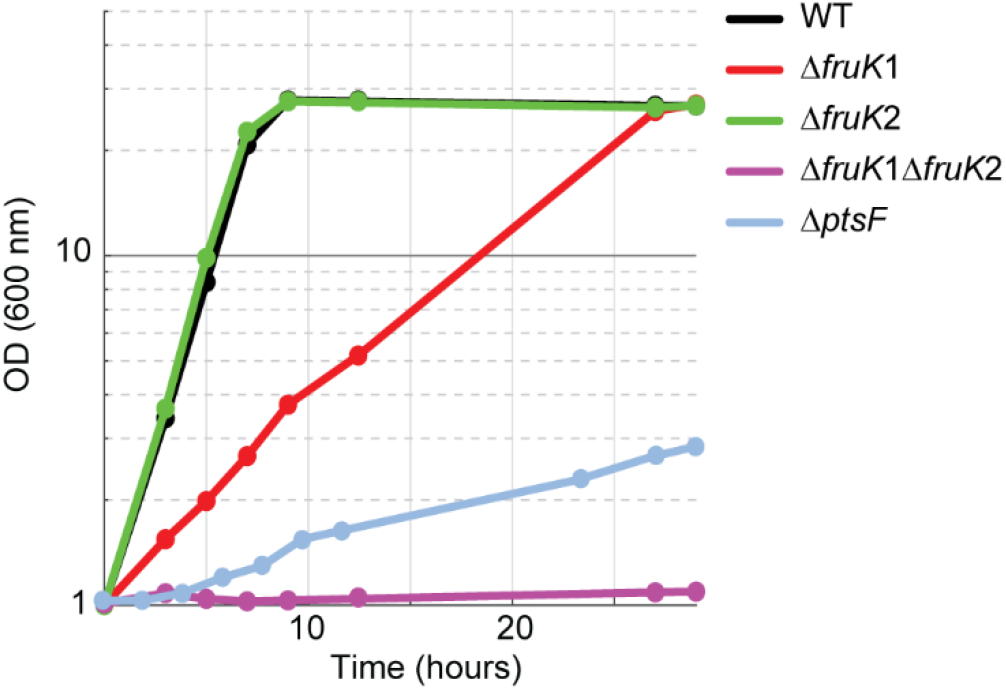
Growth on 2 % fructose of strains lacking parts of the canonical fructose utilization pathway, fructose specific PTS (*ptsF*) or 1-phosphofructokinases (*fruK1*, *fruK2*).

### Adaptive evolution for growth on fructose

To evolve the specificity of the glucose specific PTS compound towards fructose, *C. glutamicum* strains ∆*ptsF* and ∆*fruK1* ∆*fruK2* were incubated in CgXII minimal media containing 2 % fructose as a sole carbon source. After incubation for 3-4 days, all strains had grown to stationary phase. Samples from each culture were transferred to LB plates for single colony isolation. When subsequently transferred to fructose minimal medium, the isolated strains immediately showed fast growth, indicating that a mutation compensating for the growth deficiency had occurred. To increase the variance 20 cultures of each genetic backgrounds (∆*ptsF* and ∆*fruK1* ∆*fruK2*) were incubated for four days in fructose minimal medium. All strains reached stationary phase within this time. To identify if mutations in PtsG are responsible for the growth recovery, the *ptsG* locus of the isolated mutants was amplified by PCR and sequenced by Sanger-sequencing. Sequencing results revealed that all strains analyzed (n=40) had nonsynonymous substitution in the coding sequence of *ptsG*. Among these mutants, only three different point mutations were found. These mutations altered the amino acids M260V, M260T or P318S. The most abundant mutation among the three was M260V (Supplementary Figure 1).

Mutants from both the ∆*ptsF* and the ∆*fruK1* ∆*fruK2* background representing all three PtsG variants were analyzed for growth in fructose, sucrose, as well as in fructose + glucose minimal medium (Figure 3). The six analyzed mutants showed restored, fast growth with fructose as a sole source of carbon, moreover they grew to slightly higher biomass concentrations than the WT strain. In sucrose minimal medium as well as in glucose + fructose medium the strains grew similarly to the WT control and reached ODs 2-fold higher than their parental strains (∆*ptsF* or ∆*fruK1* ∆*fruK2*), since the latter can only efficiently utilize the glucose part of the provided carbon sources. While the mutant strains reached comparable maximal ODs in medium containing sucrose only, they grew to slightly higher with glucose + fructose, and to significantly higher maximal ODs when fructose was used as sole carbon source.

**Figure 3.**
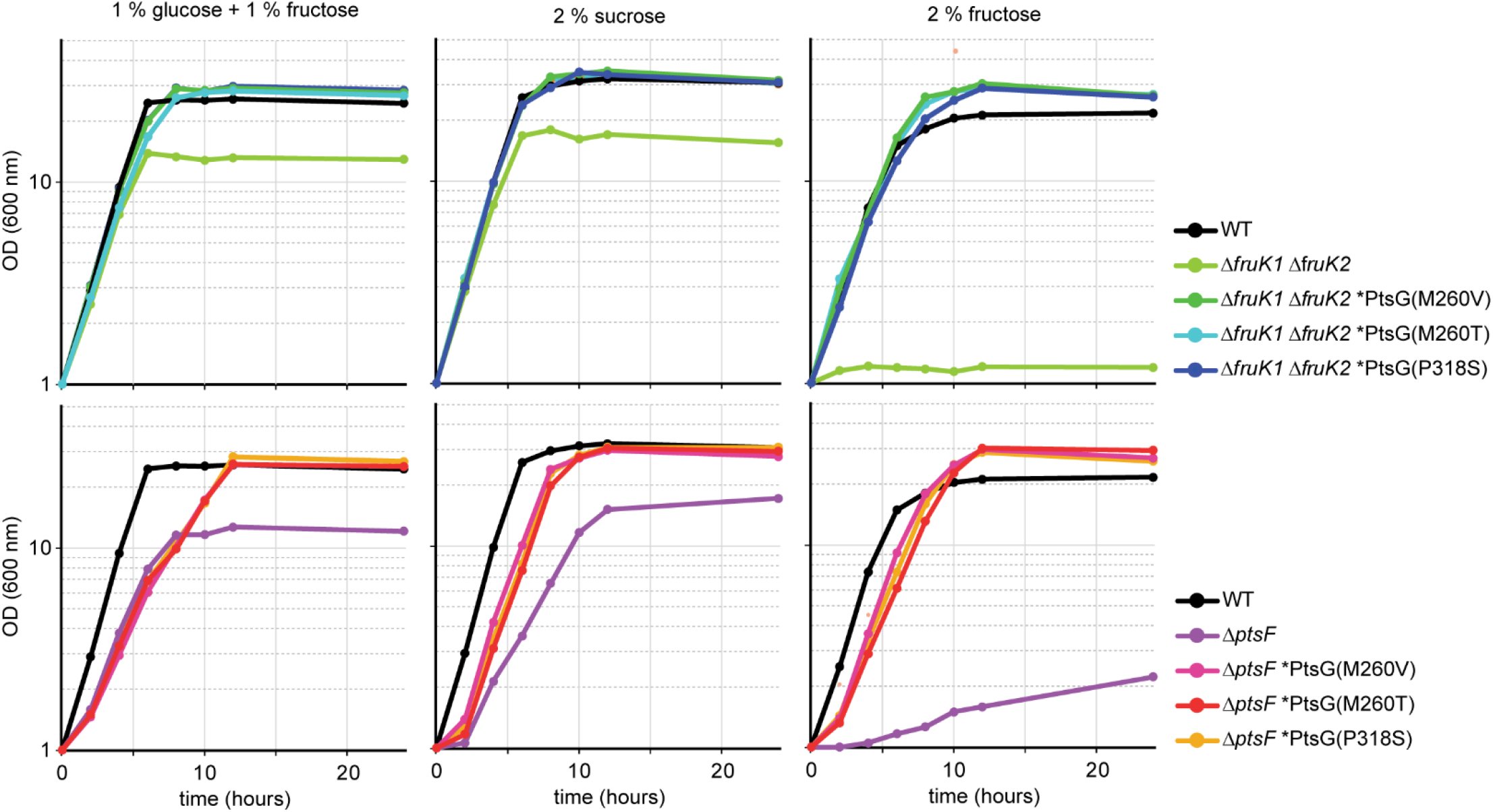
Growth of isolated fructose mutants on glucose + fructose, sucrose, and fructose.

After having shown that the mutants derived from the parental ∆*fruK1* ∆*fruK2* strain grew with fructose and contained nonsynonymous mutations in the *ptsG* locus, either the gene encoding the general PTS subunit *hpr* or the glucose specific subunit PtsG was deleted in these mutants. Both deletions *ptsG* and *hpr* in the ∆*fruK1* ∆*fruK2* mutants abolished growth with fructose in these strains (Supplementary Figure 2). Thus, the activity of the glucose specific PTS is responsible for fructose utilization in these mutants.

### Evidence for generation of fructose 6-phosphate from fructose via glucose specific PTS

To test the hypothesis that PtsG phosphorylates fructose to yield fructose 6-P, genetic experiments were performed. First, it was determined if 6-phosphofructokinase is required for fructose catabolism via glucose specific PTS. Therefore, the 6-phosphofructokinase gene (*pfkA*) was deleted in strain ∆*fruK1* ∆*fruK2,* which lacked both 1-phosphofructokinase genes, as well as in the evolved ∆*fruK1* ∆*fruK2* ALE mutant PtsG^M260T^. Both these *pfkA* deletion mutants were not able to grow in fructose minimal medium (Figure 4A), indicating that PtsG phosphorylates fructose exclusively to fructose 6-phosphate.

**Figure 4.**
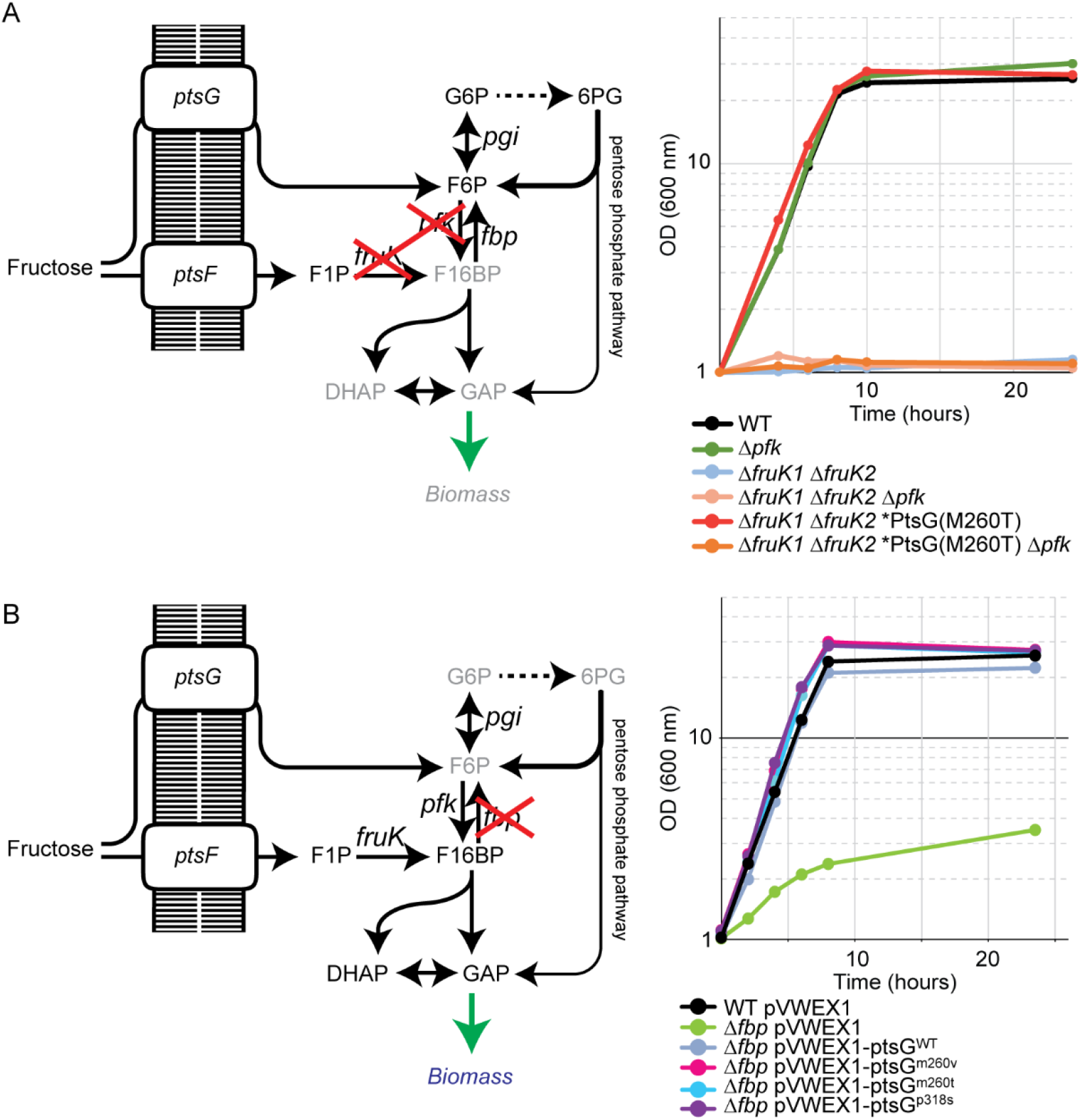
Genetic elucidation of fructose phosphorylation to fructose 6-P by PtsG. A. Deletion of 6-phosphofruktokinase (∆*pfk*) in ∆*fruK1* ∆*fruK2* fructose mutant abolishes growth. B. Overexpression of PtsG variants improve growth of a fructose 1,6-bisphosphatase deletion strain (∆*fbp*).

The deletion of *pfkA* in the WT background did not alter growth of the strain with fructose, as fructose utilized via PtsF and FruK enters glycolysis at fructose 1,6-bisphosphate. In the absence of 6-phosphofructokinase in the strain ∆*fruK1* ∆*fruK2* and the ∆*fruK1* ∆*fruK2* ALE mutant PtsG^M260T^ (see Figure 4A), the only way fructose 6-P can be catabolized is via the oxPPP. From three molecules of fructose 6-P entering the oxPPP, one molecule glyceraldehyde 3-phosphate (GAP) and three molecules of carbon dioxide are produced, while two molecules fructose 6-P are regenerated. This low feed to the “lower” metabolism seemingly is not sufficient to allow for growth. One reason might be due to sugar phosphate stress. It might also be the case that metabolism probably does not utilize the GAP produced efficiently enough as it must be used to provide PEP for fructose phosphorylation, in order to keep the stoichiometric influx of fructose 6-P.

Based on previous findings that fructose 1,6-bisphosphatase is important for fructose catabolism via PtsF and 1-phosphofructokinases FruK1 and/or FruK2 (Becker et al., 2005;Georgi et al., 2005), we hypothesized that strains growing on fructose via PtsG, synthesizing fructose 6-P directly from fructose do not require fructose 1,6-bisphosphatase for growth. To test this, we overexpressed the PtsG variants in a ∆*fbp* strain and analyzed its growth on fructose (Figure 4B). This experiment tests only for the small flux to fructose 6-P, which is necessary to generate essential PPP intermediates (erythrose 4-P, ribose 5-P, and glucose 6-P). Notably, some residual growth on fructose was observed for ∆*fbp*. This might be due to the presence of the genomic *ptsG* in this strain, which is commensurate with some flux of fructose phosphorylation in the WT as observed previously (Kiefer et al., 2004). However, all PtsG variants allowed the ∆*fbp* strain to regain growth as fast as the WT strain on fructose (Figure 4B). Thus, fructose 1,6-bisphosphatase is dispensable for growth if fructose catabolism is mediated via PtsG with fructose being directly converted to fructose 6-P.

### Complementation of ∆*ptsG* ∆*ptsF* by*ptsG* overexpression

After having shown that PtsG mediated fructose catabolism complemented for the growth impairment due to the absence of fructose 1,6-bisphosphatase, a more rigorous test was attempted. A strain lacking the genes for both fructose and glucose specific PTS subunits was constructed (∆*ptsG* ∆*ptsF*) and it was tested if PtsG and/or the selected PtsG variants support growth with fructose as sole carbon source. Strain ∆*ptsF* ∆*ptsG* revealed a clean phenotype as growth with fructose as sole carbon source was completely abolished. With glucose, however, ∆*ptsG* ∆*ptsF* showed some residual growth, which likely depended on PTS-independent glucose catabolism (Ikeda et al., 2011;Lindner et al., 2011). The observed reduction of biomass formation to 50 % with sucrose as sole carbon source reflects the fact that only the glucose moiety, but not the fructose moiety of the disaccharide can be catabolized by strain ∆*ptsF* ∆*ptsG*. Under the chosen conditions, PTS-independent fructose catabolism is irrelevant as indicated by this finding and the inability of strain ∆*ptsF* ∆*ptsG* to grow with fructose alone.

To test for *ptsG* mediated fructose utilization, strain ∆*ptsG* ∆*ptsF* was transformed with expression plasmids overexpressing either native *ptsG*, one of the three newly identified *ptsG* variants or as positive control native *ptsF*. Growth of the resulting strains in minimal media containing either fructose, glucose or sucrose is shown in Figure 5. Growth of ∆*ptsF* ∆*ptsG* with glucose was complemented to a similar extent with all PtsG variants, including native PtsG, whereas overexpression of the fructose specific PTS gene *ptsF* did not. On fructose, all constructs complemented the growth phenotype of ∆*ptsF* ∆*ptsG*, but to varying extent. Native *ptsG* supported a significantly lower growth rate (0.19 h^-1^) as compared to the growth achieved with the three mutated *ptsG*-versions (0.36 to 0.39 h^-1^) and with *ptsF* (0.44 h^-1^). Lag phases were observed only with native PtsG and with variant PtsG^P318S^ (Figure 5). Similar to the observed higher ODs reached by the ALE mutants as compared to their parent strains (Figure 3), also the reverse engineered strains analyzed grew to about 20 % higher OD in fructose minimal medium than strain ∆*ptsG* ∆*ptsF* (Table 3). With sucrose as sole carbon source, all strains grew similar to WT except for ∆*ptsF* ∆*ptsG* carrying the empty vector as negative control, which showed a lag phase, grew slower and reached about half of the OD than the other strains.

**Figure 5.**
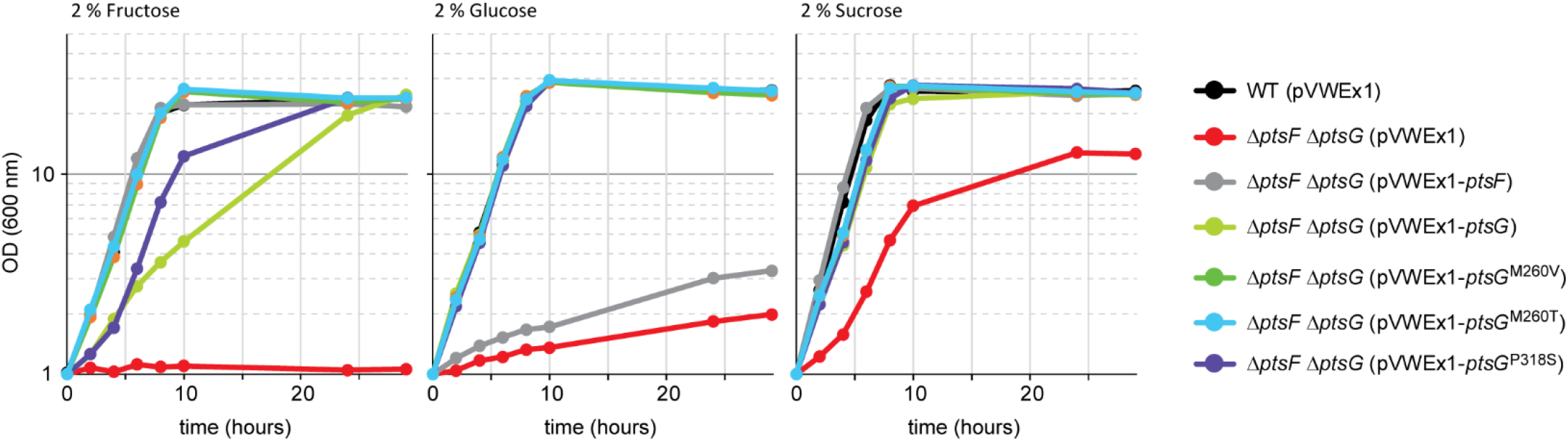
Growth complementation of the ∆*ptsF* ∆*ptsG* strain by overexpressing PtsG-variants.

**Table 3.** Growth rates and changes in OD (600 nm) during growth of recombinant strains on 2 % glucose, 2 % fructose, or 2 % sucrose.

|  | 2 % Glucose |  |  |  | 2 % Fructose |  |  |  | 2 % Sucrose |  |  |
| --- | --- | --- | --- | --- | --- | --- | --- | --- | --- | --- | --- |
| Growth rates (h<sup>-1</sup>) |  |  |  |  |  |  |  |  |  |  |  |
| WT (pVWEx1) | 0.40 | ± | 0.00 |  | 0.41 | ± | 0.01 |  | 0.49 | ± | 0.00 |
| *ΔptsF ΔptsG* (pVWEx1) | 0.03 | ± | 0.00 |  | 0 | ± | 0 |  | 0.26 | ± | 0.01 |
| *ΔptsF ΔptsG* (pVWEx1-*ptsF*) | 0.03 | ± | 0.00 |  | 0.45 | ± | 0.01 |  | 0.50 | ± | 0.01 |
| *ΔptsF ΔptsG* (pVWEx1-*ptsG*) | 0.37 | ± | 0.01 |  | 0.19 | ± | 0.01 |  | 0.33 | ± | 0.00 |
| *ΔptsF ΔptsG* (pVWEx1-*ptsG*<sup>P318S</sup>) | 0.40 | ± | 0.01 |  | 0.36 | ± | 0 |  | 0.41 | ± | 0.00 |
| *ΔptsF ΔptsG* (pVWEx1-*ptsG*<sup>M260V</sup>) | 0.40 | ± | 0.00 |  | 0.39 | ± | 0.01 |  | 0.42 | ± | 0.01 |
| *ΔptsF ΔptsG* (pVWEx1-*ptsG*<sup>M260T</sup>) | 0.40 | ± | 0.00 |  | 0.38 | ± | 0.00 |  | 0.42 | ± | 0.00 |
| ΔOD(600 nm) |  |  |  |  |  |  |  |  |  |  |  |
| WT (pVWEx1) | 27.8 | ± | 0.2 |  | 21.1 | ± | 0.2 |  | 25.6 | ± | 0.2 |
| *ΔptsF ΔptsG* (pVWEx1) | 0.8 | ± | 0.2 |  | 0.0 | ± | 0.0 |  | 11.8 | ± | 0 |
| *ΔptsF ΔptsG* (pVWEx1-*ptsF*) | 25.6 | ± | 0.1 |  | 21.7 | ± | 0.5 |  | 25.8 | ± | 1.6 |
| *ΔptsF ΔptsG* (pVWEx1-*ptsG*) | 27.7 | ± | 0.5 |  | 25.1 | ± | 0.1 |  | 24.8 | ± | 0.6 |
| *ΔptsF ΔptsG* (pVWEx1-*ptsG*<sup>P318S</sup>) | 28.1 | ± | 0.3 |  | 24.6 | ± | 0.3 |  | 26.7 | ± | 0.3 |
| *ΔptsF ΔptsG* (pVWEx1-*ptsG*<sup>M260V</sup>) | 27.7 | ± | 0.7 |  | 24.8 | ± | 0.9 |  | 27.2 | ± | 0.4 |
| *ΔptsF ΔptsG* (pVWEx1-*ptsG*<sup>M260T</sup>) | 28.4 | ± | 0 |  | 25.6 | ± | 0.6 |  | 26.8 | ± | 0.4 |

### Faster fructose uptake mediated by the selected PtsG variants

To analyze the kinetics of the PtsG mediated fructose transport we used ^14^C labeled fructose as a tracer. The kinetic data obtained from these experiments is shown in Table 4. Strains that possess PtsF showed sigmoidal dependence of the uptake rate on the fructose concentration with Hill coefficients between 2 and 3, while ∆*ptsF* mutants did not (Table 4; Supplementary Figure 3). No fructose uptake was detected by mutant ∆*ptsF* ∆*ptsG.* Fructose uptake was detected in the absence of PtsF, however, the KM value was about twenty-fold higher than the K1/2 value observed for strains that possess PtsF (Table 4). Moreover, fructose uptake was five to ten-fold faster in the presence of PtsF as compared to its absence. Thus, PtsF allowed for fast fructose uptake with high affinity, whereas PtsG supported slower uptake with lower affinity (Table 4).

**Table 4.** Kinetics parameters of fructose uptake by PtsF and PtsG.

| Strain | Hill-coefficient | Sigmoid | K<sub>1/2</sub><br>[μM] | K<sub>M</sub><br>[μM] | Vmax<br>[nmol / (min*mg(CDW))] |
| --- | --- | --- | --- | --- | --- |
| WT | 3.00 | Yes | 45.7 | - | 73.5 |
| Δ*ptsF* | / | X | - | 841 | 8.4 |
| Δ*ptsG* | 2.38 | Yes | 44.8 | - | 41.6 |
| Δ*ptsF* Δ*ptsG* | / | X | - | n.u. | - |
| Δ*ptsF* Δ*ptsG* (pVWEx1) | / | X | - | n.u. | - |
| Δ*ptsF* Δ*ptsG* (pVWEx1-*ptsF*) | 2.57 | Yes | 30.0 | - | 40.8 |
| Δ*ptsF* Δ*ptsG* (pVWEx1-*ptsG*) | / | X | - | 739 | 6.7 |
| Δ*ptsF* Δ*ptsG* (pVWEx1-*ptsG*<sup>M260V</sup>) | / | X | - | 520 | 10.0 |
| Δ*ptsF* Δ*ptsG* (pVWEx1-*ptsG*<sup>M260T</sup>) | / | X | - | 459 | 7.1 |
| Δ*ptsF* Δ*ptsG* (pVWEx1-*ptsG*<sup>P318S</sup>) | / | X | - | 325 | 12.4 |

n.u. no uptake.

The PtsG mutants showed higher affinity for fructose than WT PtsG. Graphs of fructose uptake are shown in Supplementary Figure 3. The lowest apparent K_M_ was determined for PtsG^P318S^ (325 µM), which is lower than half of WT PtsG (739 µM), but still ten-fold higher than the K1/2 value of PtsF. Two of the three PtsG variants (M260V and P318S) supported about 1.5 to 2 fold faster fructose uptake than WT PtsG (change of 6.7 to 10 and 12.4 nmol / (min*mg(CDW)), respectively). Thus, the PtsG mutations showed improved kinetic parameters for fructose uptake as compared to WT PtsG. Notably, since the maximal uptake rates observed for the PtsG mutants did not reach that supported by PtsF, their improved kinetic parameters may not explain the fast growth observed for the respective strains *in vivo*. All growth experiments performed here exceed the K_M_ concentrations by more than 100-fold, thus all PtsG variants should work under saturation conditions and affinity should not be a limiting factor.

### ^13^C labeling experiments reveal substantially higher oxPPP flux

In order to analyze altered fluxes in the strains utilizing fructose via PtsG we performed ^13^C labeling experiments. As the flux via the oxPPP and the associated NADPH provision is low during growth of *C. glutamicum* WT on fructose and depends at least in part on fructose 1,6-bisphosphatase, we hypothesized that PtsG mediated fructose catabolism directly leading to fructose 6-P instead of fructose 1,6-BP might result in a higher flux via the oxPPP. To test this hypothesis, we performed ^13^C labeling experiments with ^13^C-1-fructose as sole carbon source for growth. For comparison, cells were grown with ^13^C-1-glucose. During fructose and glucose catabolism via the oxPPP, the labeled C1 is lost as ^13^CO_2_ in the oxidative decarboxylation of gluconate 6-P to ribulose 5-P. Thus, only unlabeled ribose 5-P is present and hence histidine, which is derived from ribose 5-P is expected not to carry 13C label from the carbons derived from ribose 5-P. If the non-oxPPP is used to provide C_5_ building blocks, ^13^C-labeling of xylulose 5-P (generated by transketolase reactions) and other pentose phosphate molecules is expected. Specifically, the non-oxPPP converts two molecules of fructose 6-P (fully labeled at C1) and one molecule of GAP (50 % labeling at C3) to two molecules of xylulose 5-P (fully labeled at C1 and 50 % labeled at C3), and one molecule of unlabeled ribose 5-P as shown in detail in Supplementary Figure 4.

To obtain a clean negative control devoid of labeling patterns from the oxPPP, we included strain ∆*zwf*, which lacks glucose 6-P dehydrogenase, the entry-point of the oxPPP. Figure 6 shows the labeling patterns of histidine and alanine in the ^13^C labeling experiments performed with *C. glutamicum* WT and the indicated mutants. Our results confirmed previous findings that the oxPPP flux is lower during growth with fructose than during growth with glucose (Kiefer et al., 2004) since labeling in L-alanine and L-histidine were higher with ^13^C-1-fructose than with ^13^C-1-glucose. In the WT oxPPP-flux is barely active during growth on fructose, indicated by the high ^13^C-labeling in L-alanine and L-histidine during growth with ^13^C-1-fructose, which was almost as high as in the ∆*zwf* strain, which lacks the oxPPP. The ∆*ptsF* ∆*ptsG* strain expressing *ptsF* showed a labeling pattern similar to WT, which indicated a low ox-PPP flux when fructose was catabolized via PtsF. On the other hand, ∆*ptsF* ∆*ptsG* strains overexpressing the PtsG variants showed reduced absolute labeling of L-alanine and L-histidine. This provided evidence for a higher oxPPP flux when fructose is utilized via PtsG with direct conversion to fructose 6-P. Notably, the observed labeling is similar to the labeling observed when WT and the other strains grew on ^13^C-1-glucose (Figure 6 and data not shown). The absolute labeling in L-alanine and L-histidine (^13^C-abundance reduced by about 20 % in ∆*ptsF* ∆*ptsG* strains overexpressing the PtsG variants compared to the ∆*zwf* strain used as reference without oxPPP flux) allowed to calculate that about 20 % of the fructose molecules were catabolized via the oxPPP, which was comparable to the oxPPP flux observed during growth with glucose.

**Figure 6.**
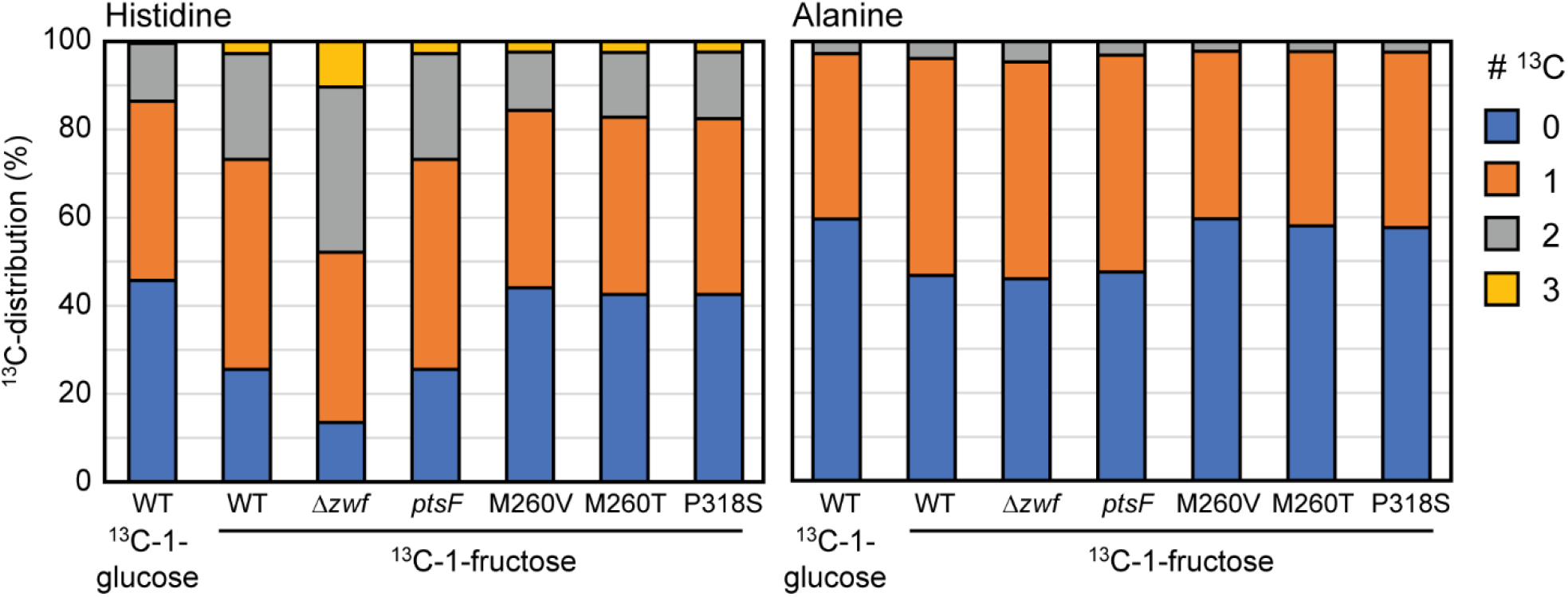
^13^C-labeling in histidine and alanine in WT, ∆*zwf* and the ∆*ptsF* ∆*ptsG* strain overexpressing *ptsF* or the *ptsG* variants upon feeding ^13^C-1-fructose or ^13^C-1-glucose.

### Evolved and reverse engineered strains showed reduced over flow metabolism

Fast growth with glucose is known to be associated with intermittent lactate accumulation in the culture medium. During the exponential growth phase NAD-dependent L-lactate dehydrogenase reduced pyruvate to L-lactate, which is secreted (Dominguez et al., 1998). L-lactate is re-utilized after induction of LldR by L-lactate and derepression of the *lld* operon for L-lactate catabolism (Georgi et al., 2008). The observed higher oxPPP flux when fructose is catabolized via PtsG as compared to PtsF prompted us to investigate metabolic consequences. As already described above, the isolated mutants as well as the reverse engineered strains grew to higher biomass concentrations than the *C. glutamicum* WT. Here, we investigated whether metabolic consequences can be observed with regard to byproduct formation. Besides growing to higher ODs, the mutant and the reverse engineered strain accumulated less L-lactate during growth (Figure 7).

**Figure 7.**
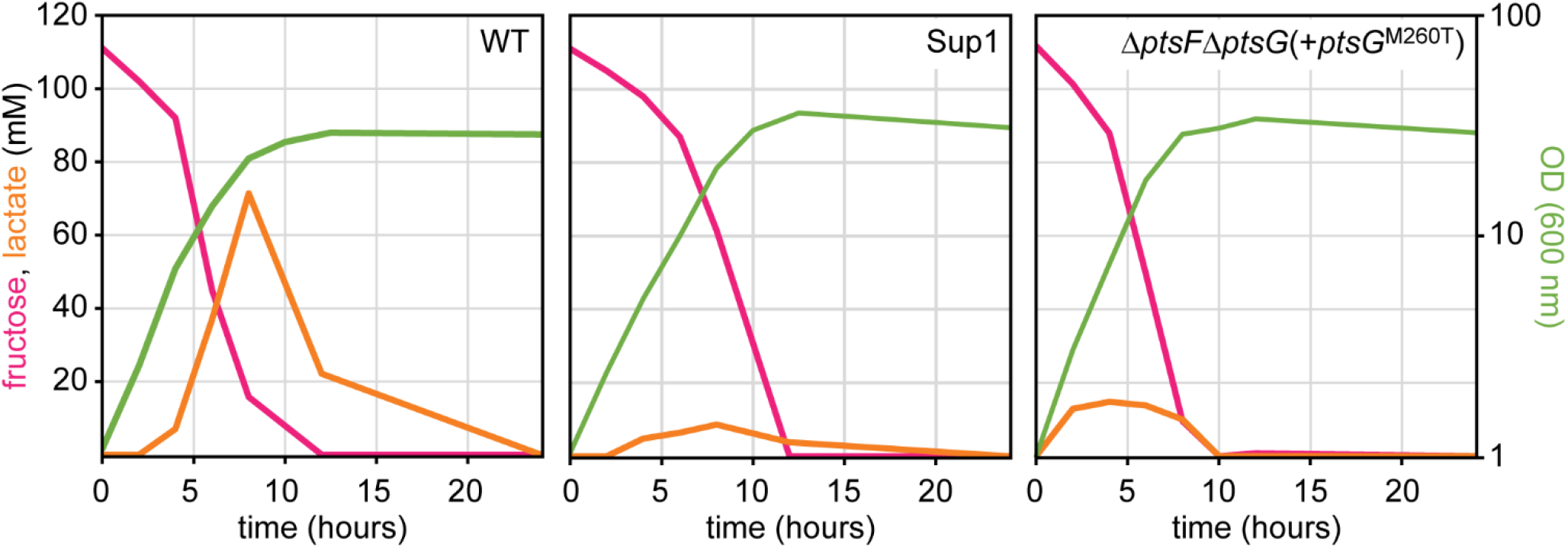
Transient lactate accumulation in cultures of WT, mutant and reconstructed strain during growth on fructose.

### PtsG catalyzed fructose utilization improved L-lysine production

L-lysine production benefits from improved NADPH provision and reduced by-product formation. Since we have shown that PtsG mediated fructose catabolism is characterized by increased oxPPP flux relevant for NADPH provision as well as reduced intermittent formation of L-lactate as by-product, the metabolic consequence on L-lysine production was determined. First low-level L-lysine production by the ALE mutants was enabled via transformation using a plasmid for overexpression of a feedback resistant aspartate kinase gene (pVWEx1-*lysC^fbr^*). Compared to the control, i.e., *C. glutamicum* WT(pVWEx1-*lysC^fbr^*) L-lysine production in minimal medium containing 40 g/L fructose was increased about 5-fold. The ALE mutants overexpressing *lysC^fbr^* produced about 26 mM, while the WT overexpressing *lysC^fbr^* produced 4.5 mM.

After these initial and very promising results, we constructed the PtsG pathway as sole route for fructose utilization in the L-lysine producer CgLYS4 ∆*ptsF* (Sgobba et al., 2018). This strain is a *ptsF* deletion strain derived from CgLYS4, thus, lacks native fructose utilization via PtsF. CgLYS4 produces L-lysine due to feedback resistant aspartokinase, attenuated homoserine dehydrogenase and improved pyruvate carboxylase (*lysC^T311I^, hom^V59A^, pyc^P458S^*) and the strain shows reduced by-product formation as it carries deletions in the lactate dehydrogenase gene (*ldhA*) and acetate production genes (*pta*-*ackA*). Production of L-lysine was analyzed after growth with 40 g/L fructose as sole carbon source and compared to the production achieved with the parental strain CgLYS4 (Sgobba et al., 2018). All tested strains utilizing fructose via PtsG accumulated about 50% more L-lysine than the PtsF-positive CgLYS4 (about 45 mM as compared to about 30 mM; Table 5). Notably, the strain overexpressing native PtsG reached a similar final L-lysine concentration as the strains overexpressing the new PtsG variants, but much later since this strain grew significantly slower (data not shown). Interestingly, overexpression of *ptsF* reduced L-lysine production compared to CgLYS4. Taken together fructose catabolism via the isolated PtsG variants is a promising strategy to improve L-lysine production.

**Table 5.** L-lysine production of strains utilizing fructose via PtsG. Data represents means of three independent experiment with errors < 5 %.

| Strain | L-lysine (mM) |
| --- | --- |
| CgLYS4 | 29.7 |
| CgLYS4 $\Delta ptsF$ pVWEx1-*ptsG* | 43.1 |
| CgLYS4 $\Delta ptsF$ pVWEx1-*ptsG*<sup>P318S</sup> | 44.2 |
| CgLYS4 $\Delta ptsF$ pVWEx1-*ptsG*<sup>M260V</sup> | 46.9 |
| CgLYS4 $\Delta ptsF$ pVWEx1-*ptsG*<sup>M260T</sup> | 46.2 |
| CgLYS4 $\Delta ptsF$ pVWEx1-*ptsF* | 24.0 |

## Discussion

In this study, ALE was used to isolate mutants able to catabolize fructose via PtsG. The PtsG variants enabled fast growth with fructose with increased oxPPP flux. Production of L-lysine was chosen as an application example and improved L-lysine titers associated with fast growth on fructose.

As shown here, only a few days of cultivation under selective conditions were sufficient to achieve the desired growth phenotype indicating that mutation of a single gene was sufficient and that several mutations in this gene resulted in the desired growth phenotype. This is not unprecedented as earlier studies revealed the ability of *C. glutamicum* to evolve relatively quickly into a niche or to overcome a genetic impairment (Youn et al., 2009;Lindner et al., 2011;Uhde et al., 2013). The present and these studies share that they selected for utilization of a carbon and energy source. In addition, ALE has been used to select *C. glutamicum* mutants withstanding adverse conditions e.g. due to methanol or indole (Lessmeier and Wendisch, 2015;Hennig et al., 2020;Kuepper et al., 2020;Walter et al., 2020) or mutants that have overcome the requirement for an additive such as iron chelator PCA (Graf et al., 2019) or production of. e.g., putrescine (Jorge et al., 2017;Li and Liu, 2017). Thus, ALE has proven valuable for *C. glutamicum* metabolic engineering (Stella et al., 2019).

Apart from *C. glutamicum*’s PtsG also the mannose PTS system of *E. coli* generates fructose 6-P from fructose (Kornberg, 2001), this tendency of promiscuity of the PTS system compounds also applies to some sugar kinases, e.g. *E. coli’s* enzymes xylulokinase is active on xylulose and ribulose (Di Luccio et al., 2007) and fuculokinase is active on fuculose and ribulose (LeBlanc and Mortlock, 1971). One explanation for their promiscuity is that carbohydrate kinases are ancient enzymes, which needed to evolve into niches of present carbon sources (Roy et al., 2019). A good example for enzyme promiscuity is the fast evolution for utilization of new substrates shown for *E. coli* (Guzman et al., 2019). Regarding PTS specificity, a prominent and promiscuous example is probably *E. coli*’s mannose PTS compound, which besides mannose also takes glucose, fructose, N-acetylglucosamine and glucosamine (Curtis and Epstein, 1975;Chou et al., 1994). Similar to the approach described here, Wang *et al.* used a *C. glutamicum* ∆*ptsF* strain, evolved it on sucrose and found suppressor mutants with inactivated 1-phosphofructokinase gene, indicating the role of sugar-phosphates in transcriptional repression, likely of *ptsG*, which might explain the enhanced NADPH and L-lysine production from sucrose and fructose (Wang et al., 2016). The responsible regulator of sugar utilization, SugR, represses expression of *ptsG* (Engels and Wendisch, 2007), deletion of *sugR* derepressed *ptsG* transcription and consequently facilitates glucose utilization and improved L-lysine productivity (Perez-Garcia et al., 2016).

The results deduced from the ^13^C-labeling obtained in our study, show some differences to the data described by Kiefer et al., that might be explained by the use of a L-lysine producer strain in the study by Kiefer et al. since L-lysine overproduction provides a strong NADPH sink. *C. glutamicum* can respond to different metabolic burdens differing in their NADPH requirements, as was shown in a metabolic flux comparison of *C. glutamicum* WT grown either under standard conditions or upon triggering L-glutamate production and of a L-lysine producing strain (Marx 1997). Flux in the oxPPP and, thus, NADPH generation was highest in the lysine producer, intermediate in WT and lowest under L-glutamate production (isocitrate dehydrogenase provides NADPH and 2-oxoglutarate to balance the NADPH requirement of glutamate dehydrogenase for reductive amination of 2-oxoglutarate to yield glutamate (Bormann et al., 1992).

We used L-lysine production as a readout to prove the increased NADPH-availability in the engineered strains. In addition to its NADPH demand, L-lysine production highly depends on strong fluxes towards anaplerosis, providing oxaloacetate as the precursor for aspartate biosynthesis, the starting point of L-lysine biosynthesis. The supply anaplerotic precursors might be negatively affected by the PEP dependent sugar phosphorylation carried out by the PTS system, as PTS independent sugar utilizations improved L-lysine production (Lindner et al., 2011). However, using ATP dependent sugar phosphorylation, e.g. fructokinase was shown to have a negative effect on ATP availability, and hence sugar uptake (Xu et al., 2020). Recently for L-lysine and L-threonine production (both high-NADPH demanding products) optimal flux ratio between oxPPP and glycolysis was determined (Murai et al., 2020), indicating a high demand of oxPPP for these products. Similar to the effects seen for L-lysine, our discovery might be of value for biotechnological use for high-NADPH-depending products, e.g. threonine or 1,5-diaminopentane. This approach may be paired with others: Further approaches tackling NADPH recovery for increased bioproductions are, overexpression of membrane bound transhydrogenase (Kabus et al., 2007), deletion of phosphoglucose isomerase (Marx et al., 2003), overexpression of NAD kinase (Lindner et al., 2010), overexpression of oxPPP enzymes (Becker et al., 2007). However, exclusively focusing on glucose as a carbon source. Indirect effects of by similarly to the approach described above were undertaken by overexpressing gluconeogenetic fructose 1,6-bisphosphatase, which increased L-lysine production from fructose (Georgi et al., 2005).

A new and fast way of fructose utilization via the optimized PtsG variants was shown in the biotechnologically important microbe *C. glutamicum*. The pathway allows an increased oxPPP flux and consequently an improved NADPH regeneration rate, which was exploited here for the high NADPH demanding L-lysine production. The here described results and especially the PtsG mutations might also be advantageous for the production of other NADPH demanding products, e.g. other amino acids or derivatives like diamines, and fatty acids.

## Acknowledgements

We thank Änne Michaelis for assistance with LC-MS analysis of ^13^C-labeling in amino acids.

## Author Contributions

S.N.L. and V.F.W. conceived the study. S.N.L. and V.F.W. designed the experiments. I.K., D.B., J.P.K., T.M., and S.N.L. performed metabolic engineering experiments. N.R. and G.M.S. performed sugar uptake experiments. S.N.L. and V.F.W. analyzed the results. S.N.L., V.F.W. and G.M.S. wrote the manuscript with contributions from all authors. All authors agreed with the final version of the manuscript.

## Competing Financial Interests

The authors declare no competing interests.

## Notes

### Competing Interest Statement

The authors have declared no competing interest.

## References

Becker, J., Klopprogge, C., Herold, A., Zelder, O., Bolten, C.J., and Wittmann, C. (2007). Metabolic flux engineering of L-lysine production in *Corynebacterium glutamicum*--over expression and modification of G6P dehydrogenase. J Biotechnol 132, 99–109.

Becker, J., Klopprogge, C., Zelder, O., Heinzle, E., and Wittmann, C. (2005). Amplified expression of fructose 1,6-bisphosphatase in *Corynebacterium glutamicum* increases in vivo flux through the pentose phosphate pathway and lysine production on different carbon sources. Appl Environ Microbiol 71, 8587–8596.

Becker, J., Rohles, C.M., and Wittmann, C. (2018). Metabolically engineered *Corynebacterium glutamicum* for bio-based production of chemicals, fuels, materials, and healthcare products. Metab Eng 50, 122–141.

Blombach, B., Riester, T., Wieschalka, S., Ziert, C., Youn, J.W., Wendisch, V.F., and Eikmanns, B.J. (2011). *Corynebacterium glutamicum* tailored for efficient isobutanol production. Appl Environ Microbiol 77, 3300–3310.

Bormann, E.R., Eikmanns, B.J., and Sahm, H. (1992). Molecular analysis of the *Corynebacterium glutamicum gdh* gene encoding glutamate dehydrogenase. Mol Microbiol 6, 317–326.

Chou, C.H., Bennett, G.N., and San, K.Y. (1994). Effect of modulated glucose uptake on high-level recombinant protein production in a dense Escherichia coli culture. Biotechnol Prog 10, 644–647.

Curtis, S.J., and Epstein, W. (1975). Phosphorylation of D-glucose in *Escherichia coli* mutants defective in glucosephosphotransferase, mannosephosphotransferase, and glucokinase. J Bacteriol 122, 1189–1199.

Di Luccio, E., Petschacher, B., Voegtli, J., Chou, H.T., Stahlberg, H., Nidetzky, B., and Wilson, D.K. (2007). Structural and kinetic studies of induced fit in xylulose kinase from Escherichia coli. J Mol Biol 365, 783–798.

Dominguez, H., and Lindley, N.D. (1996). Complete Sucrose Metabolism Requires Fructose Phosphotransferase Activity in *Corynebacterium glutamicum* To Ensure Phosphorylation of Liberated Fructose. Appl Environ Microbiol 62, 3878–3880.

Dominguez, H., Rollin, C., Guyonvarch, A., Guerquin-Kern, J.L., Cocaign-Bousquet, M., and Lindley, N.D. (1998). Carbon-flux distribution in the central metabolic pathways of *Corynebacterium glutamicum* during growth on fructose. Eur J Biochem 254, 96–102.

Eggeling, L., and Bott, M. (2005). Handbook of Corynebacterium glutamicum. CRC press.

Engels, V., and Wendisch, V.F. (2007). The DeoR-type regulator SugR represses expression of *ptsG* in *Corynebacterium glutamicum*. J Bacteriol 189, 2955–2966.

Erb, T.J., Jones, P.R., and Bar-Even, A. (2017). Synthetic metabolism: metabolic engineering meets enzyme design. Current Opinion in Chemical Biology 37, 56–62.

Georgi, T., Engels, V., and Wendisch, V.F. (2008). Regulation of L-lactate utilization by the FadR-type regulator LldR of *Corynebacterium glutamicum*. J Bacteriol 190, 963–971.

Georgi, T., Rittmann, D., and Wendisch, V.F. (2005). Lysine and glutamate production by *Corynebacterium glutamicum* on glucose, fructose and sucrose: roles of malic enzyme and fructose-1,6-bisphosphatase. Metab Eng 7, 291–301.

Giavalisco, P., Li, Y., Matthes, A., Eckhardt, A., Hubberten, H.M., Hesse, H., Segu, S., Hummel, J., Köhl, K., and Willmitzer, L. (2011). Elemental formula annotation of polar and lipophilic metabolites using 13C, 15N and 34S isotope labelling, in combination with high‐resolution mass spectrometry. Plant J 68, 364–376.

Graf, M., Haas, T., Muller, F., Buchmann, A., Harm-Bekbenbetova, J., Freund, A., Niess, A., Persicke, M., Kalinowski, J., Blombach, B., and Takors, R. (2019). Continuous Adaptive Evolution of a Fast-Growing *Corynebacterium glutamicum* Strain Independent of Protocatechuate. Front Microbiol 10, 1648.

Guzman, G.I., Sandberg, T.E., Lacroix, R.A., Nyerges, A., Papp, H., De Raad, M., King, Z.A., Hefner, Y., Northen, T.R., Notebaart, R.A., Pal, C., Palsson, B.O., Papp, B., and Feist, A.M. (2019). Enzyme promiscuity shapes adaptation to novel growth substrates. Mol Syst Biol 15, e8462.

Hennig, G., Haupka, C., Brito, L.F., Ruckert, C., Cahoreau, E., Heux, S., and Wendisch, V.F. (2020). Methanol-Essential Growth of *Corynebacterium glutamicum*: Adaptive Laboratory Evolution Overcomes Limitation due to Methanethiol Assimilation Pathway. Int J Mol Sci 21.

Ikeda, M. (2012). Sugar transport systems in *Corynebacterium glutamicum*: features and applications to strain development. Appl Microbiol Biotechnol 96, 1191–1200.

Ikeda, M., Mizuno, Y., Awane, S., Hayashi, M., Mitsuhashi, S., and Takeno, S. (2011). Identification and application of a different glucose uptake system that functions as an alternative to the phosphotransferase system in *Corynebacterium glutamicum*. Appl Microbiol Biotechnol 90, 1443–1451.

Jorge, J.M., Nguyen, A.Q., Perez-Garcia, F., Kind, S., and Wendisch, V.F. (2017). Improved fermentative production of gamma-aminobutyric acid via the putrescine route: Systems metabolic engineering for production from glucose, amino sugars, and xylose. Biotechnol Bioeng 114, 862–873.

Kabus, A., Georgi, T., Wendisch, V.F., and Bott, M. (2007). Expression of the Escherichia coli pntAB genes encoding a membrane-bound transhydrogenase in Corynebacterium glutamicum improves L-lysine formation. Appl Microbiol Biotechnol 75, 47–53.

Kiefer, P., Heinzle, E., Zelder, O., and Wittmann, C. (2004). Comparative metabolic flux analysis of lysine-producing *Corynebacterium glutamicum* cultured on glucose or fructose. Appl Environ Microbiol 70, 229–239.

Kornberg, H.L. (2001). Routes for fructose utilization by *Escherichia coli*. J Mol Microbiol Biotechnol 3, 355–359.

Kuepper, J., Otto, M., Dickler, J., Behnken, S., Magnus, J., Jager, G., Blank, L.M., and Wierckx, N. (2020). Adaptive laboratory evolution of *Pseudomonas putida* and *Corynebacterium glutamicum* to enhance anthranilate tolerance. Microbiology (Reading) 166, 1025–1037.

Leblanc, D.J., and Mortlock, R.P. (1971). Metabolism of D-arabinose: origin of a D-ribulokinase activity in *Escherichia coli*. J Bacteriol 106, 82–89.

Lessmeier, L., and Wendisch, V.F. (2015). Identification of two mutations increasing the methanol tolerance of *Corynebacterium glutamicum*. BMC Microbiol 15, 216.

Li, Z., and Liu, J.Z. (2017). Transcriptomic Changes in Response to Putrescine Production in Metabolically Engineered *Corynebacterium glutamicum*. Front Microbiol 8, 1987.

Lindner, S.N., Niederholtmeyer, H., Schmitz, K., Schoberth, S.M., and Wendisch, V.F. (2010). Polyphosphate/ATP-dependent NAD kinase of *Corynebacterium glutamicum*: biochemical properties and impact of *ppnK* overexpression on lysine production. Appl Microbiol Biotechnol 87, 583–593.

Lindner, S.N., Seibold, G.M., Henrich, A., Kramer, R., and Wendisch, V.F. (2011). Phosphotransferase system-independent glucose utilization in *Corynebacterium glutamicum* by inositol permeases and glucokinases. Appl Environ Microbiol 77, 3571–3581.

Marx, A., De Graaf, A.A., Wiechert, W., Eggeling, L., and Sahm, H. (1996). Determination of the fluxes in the central metabolism of *Corynebacterium glutamicum* by nuclear magnetic resonance spectroscopy combined with metabolite balancing. Biotechnol Bioeng 49, 111–129.

Marx, A., Hans, S., Mockel, B., Bathe, B., De Graaf, A.A., Mccormack, A.C., Stapleton, C., Burke, K., O'donohue, M., and Dunican, L.K. (2003). Metabolic phenotype of phosphoglucose isomerase mutants of *Corynebacterium glutamicum*. J Biotechnol 104, 185–197.

Mindt, M., Walter, T., Kugler, P., and Wendisch, V.F. (2020). Microbial Engineering for Production of N-Functionalized Amino Acids and Amines. Biotechnol J 15, e1900451.

Moon, M.W., Park, S.Y., Choi, S.K., and Lee, J.K. (2007). The phosphotransferase system of *Corynebacterium glutamicum*: features of sugar transport and carbon regulation. J Mol Microbiol Biotechnol 12, 43–50.

Murai, K., Sasaki, D., Kobayashi, S., Yamaguchi, A., Uchikura, H., Shirai, T., Sasaki, K., Kondo, A., and Tsuge, Y. (2020). Optimal Ratio of Carbon Flux between Glycolysis and the Pentose Phosphate Pathway for Amino Acid Accumulation in *Corynebacterium glutamicum*. ACS Synth Biol 9, 1615–1622.

Parche, S., Burkovski, A., Sprenger, G.A., Weil, B., Kramer, R., and Titgemeyer, F. (2001). Corynebacterium glutamicum: a dissection of the PTS. J Mol Microbiol Biotechnol 3, 423–428.

Perez-Garcia, F., Peters-Wendisch, P., and Wendisch, V.F. (2016). Engineering *Corynebacterium glutamicum* for fast production of L-lysine and L-pipecolic acid. Appl Microbiol Biotechnol 100, 8075–8090.

Peters-Wendisch, P.G., Schiel, B., Wendisch, V.F., Katsoulidis, E., Mockel, B., Sahm, H., and Eikmanns, B.J. (2001). Pyruvate carboxylase is a major bottleneck for glutamate and lysine production by *Corynebacterium glutamicum*. J Mol Microbiol Biotechnol 3, 295–300.

Radek, A., Muller, M.F., Gatgens, J., Eggeling, L., Krumbach, K., Marienhagen, J., and Noack, S. (2016). Formation of xylitol and xylitol-5-phosphate and its impact on growth of d-xylose-utilizing *Corynebacterium glutamicum* strains. J Biotechnol 231, 160–166.

Rittmann, D., Lindner, S.N., and Wendisch, V.F. (2008). Engineering of a glycerol utilization pathway for amino acid production by *Corynebacterium glutamicum*. Applied and environmental microbiology 74, 6216–6222.

Rittmann, D., Schaffer, S., Wendisch, V.F., and Sahm, H. (2003). Fructose-1,6-bisphosphatase from *Corynebacterium glutamicum*: expression and deletion of the *fbp* gene and biochemical characterization of the enzyme. Arch Microbiol 180, 285–292.

Roy, S., Vivoli Vega, M., and Harmer, N.J. (2019). Carbohydrate Kinases: A Conserved Mechanism Across Differing Folds. Catalysts 9, 29.

Schafer, A., Tauch, A., Jager, W., Kalinowski, J., Thierbach, G., and Puhler, A. (1994). Small mobilizable multi-purpose cloning vectors derived from the *Escherichia coli* plasmids pK18 and pK19: selection of defined deletions in the chromosome of *Corynebacterium glutamicum*. Gene 145, 69–73.

Sgobba, E., Stumpf, A.K., Vortmann, M., Jagmann, N., Krehenbrink, M., Dirks-Hofmeister, M.E., Moerschbacher, B., Philipp, B., and Wendisch, V.F. (2018). Synthetic *Escherichia coli*-*Corynebacterium glutamicum* consortia for L-lysine production from starch and sucrose. Bioresour Technol 260, 302–310.

Stella, R.G., Wiechert, J., Noack, S., and Frunzke, J. (2019). Evolutionary engineering of *Corynebacterium glutamicum*. Biotechnol J 14, e1800444.

Uhde, A., Youn, J.W., Maeda, T., Clermont, L., Matano, C., Kramer, R., Wendisch, V.F., Seibold, G.M., and Marin, K. (2013). Glucosamine as carbon source for amino acid-producing *Corynebacterium glutamicum*. Appl Microbiol Biotechnol 97, 1679–1687.

Walter, T., Al Medani, N., Burgardt, A., Cankar, K., Ferrer, L., Kerbs, A., Lee, J.H., Mindt, M., Risse, J.M., and Wendisch, V.F. (2020). Fermentative N-Methylanthranilate Production by Engineered *Corynebacterium glutamicum*. Microorganisms 8.

Wang, Z., Chan, S.H.J., Sudarsan, S., Blank, L.M., Jensen, P.R., and Solem, C. (2016). Elucidation of the regulatory role of the fructose operon reveals a novel target for enhancing the NADPH supply in *Corynebacterium glutamicum*. Metab Eng 38, 344–357.

Wendisch, V.F. (2020). Metabolic engineering advances and prospects for amino acid production. Metab Eng 58, 17–34.

Xu, J., Zhang, J., Guo, Y., Zai, Y., and Zhang, W. (2013). Improvement of cell growth and L-lysine production by genetically modified *Corynebacterium glutamicum* during growth on molasses. J Ind Microbiol Biotechnol 40, 1423–1432.

Xu, J.Z., Ruan, H.Z., Yu, H.B., Liu, L.M., and Zhang, W. (2020). Metabolic engineering of carbohydrate metabolism systems in *Corynebacterium glutamicum* for improving the efficiency of L-lysine production from mixed sugar. Microb Cell Fact 19, 39.

Youn, J.W., Jolkver, E., Kramer, R., Marin, K., and Wendisch, V.F. (2009). Characterization of the dicarboxylate transporter DctA in *Corynebacterium glutamicum*. J Bacteriol 191, 5480–5488.

Zhao, N., Qian, L., Luo, G., and Zheng, S. (2018). Synthetic biology approaches to access renewable carbon source utilization in *Corynebacterium glutamicum*. Appl Microbiol Biotechnol 102, 9517–9529.

