## Supplementary Material for "Evolving a new efficient mode of fructose utilization for improved bioproduction in *Corynebacterium glutamicum*"

***Supplementary Material to***

**Krahn *et al.***

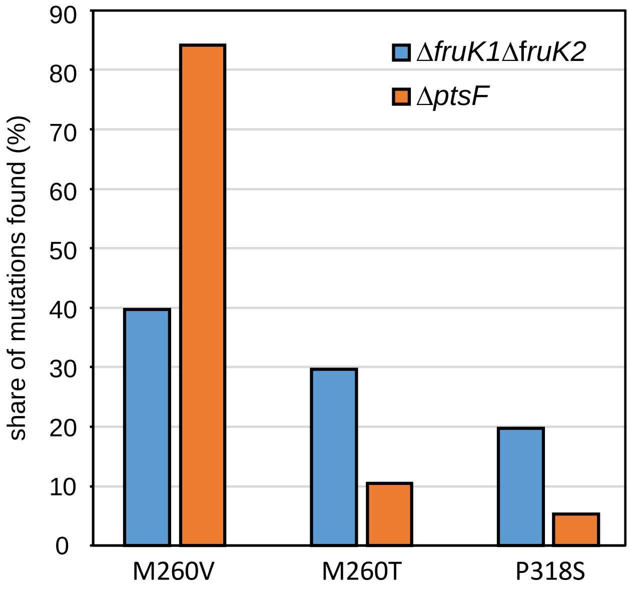


**Supplementary Figure 1.** Distribution of mutations found in *ptsG* in the mutants of Δ*fruK1* Δ*fruK2* and Δ*ptsF*, isolated after restored growth in fructose minimal medium.

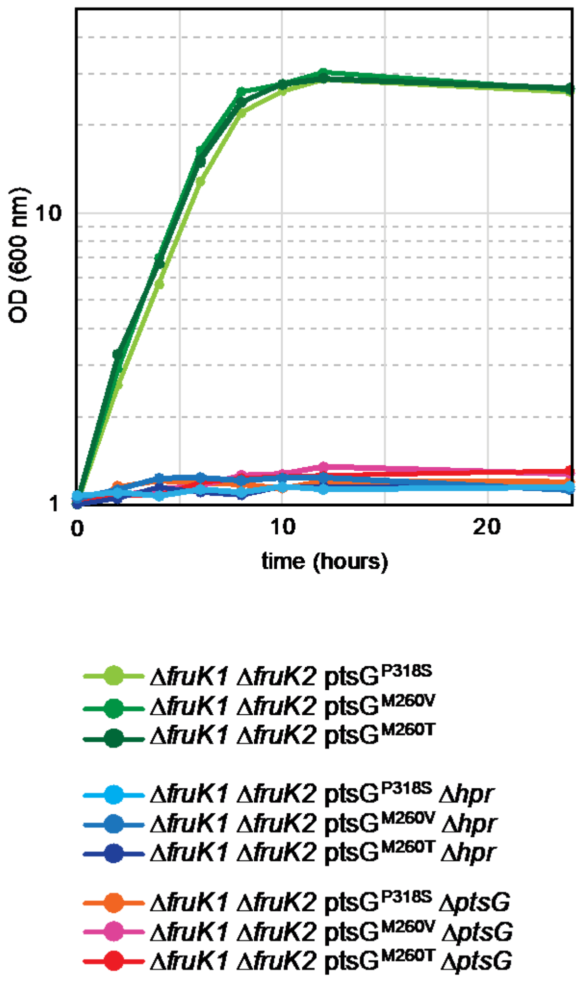


**Supplementary Figure 2.** Deletion of PTS-components *ptsG* and *hpr* abolishes growth in Δ*fruK1* Δ*fruK2* derived fructose mutants and reveals fructose utilization via PtsG.

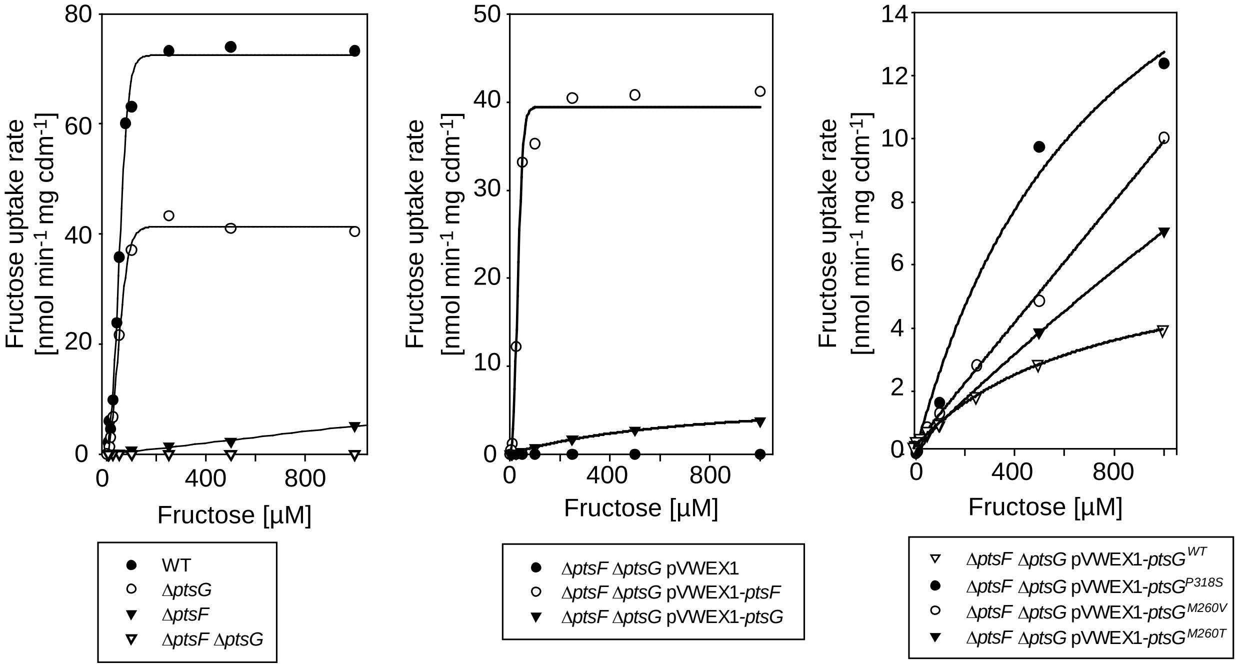


**Supplementary Figure 3:** Kinetics of fructose uptake via PtsF and PtsG WT and mutated variants of PtsG. Data represents mean values +/- SD from three independent experiments (n = 3).

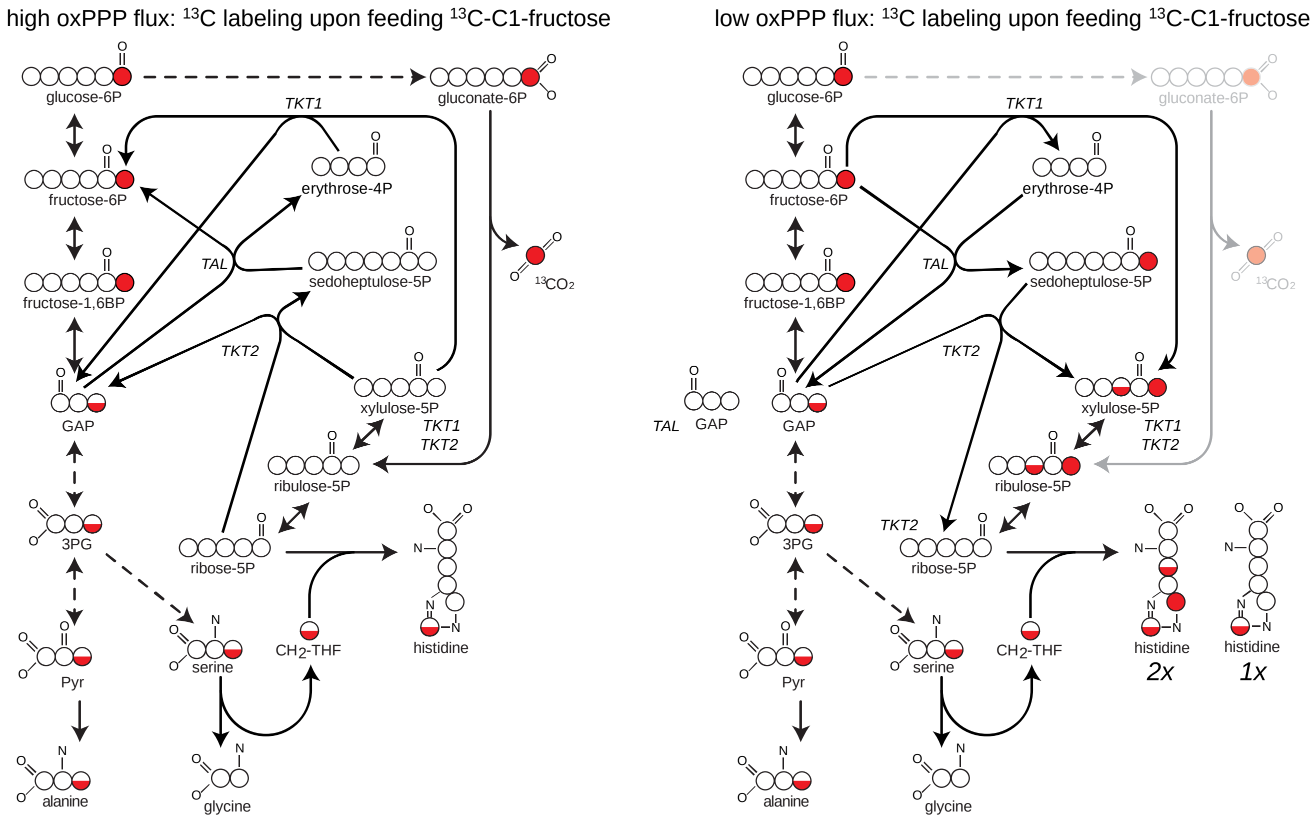


**Supplementary Figure 4.** Expected metabolic distribution of $^{13}$C-labeling from $^{13}$C-1-fructose or $^{13}$C-1-glucose assuming high or low oxidative PPP flux.

**Supplementary Table 1.** Oligonucleotide primers used in this study.

| name | Sequence (5' → 3') | purpose |
|---|---|---|
| ptsG_up | GCAGCTCGCTGCTCTTC | Amplification of *ptsG* locus - Sequencing of *ptsG* |
| ptsG_down | GATGTCTTGGCCAAAAGCTTC | Amplification of *ptsG* locus - Sequencing of *ptsG* |
| ptsG-Seq1 | GTGCGGGCATAATCTGACAG | Sequencing of *ptsG* |
| ptsG-Seq2 | GCTACAGAGAGTTCATCCAAGAAG | Sequencing of *ptsG* |
| ptsG-Seq3 | GGTCTTCTACTTCCTGCCAATTATG | Sequencing of *ptsG* |
| ptsG-Seq4 | CTGATTATGATCCCAGCGACC | Sequencing of *ptsG* |
| ptsG-Seq5 | GTTTGCTCGGCGGCATTTC | Sequencing of *ptsG* |
| ptsG-Seq6 | GAAGGCAGAAGCTAATGCAACTC | Sequencing of *ptsG* |
| ptsG-Seq7 | GAAACACCGTTGTTGCTCCAG | Sequencing of *ptsG* |
| ptsG_fw | CGTCTAGAGAAAGGAGGCCCTTCAGATGGCGTCCAAACTGACGA | Amplification of *ptsG* for cloning via *Xba*I |
| ptsG-rv | TCTAGATTACTCGTTCTTGCCGTT | Amplification of *ptsG* for cloning via *Xba*I |
| ptsF_rv | GATCTAGATTATGCGTTTACAGCTGCTTGTTG | Amplification of *ptsF* for cloning via *Xba*I |
| ptsF_fw | GGGTCTAGAGAAAGGAGGCCCTTCAGATGAATAGCGTAAATAATTCCTCGCTTG | Amplification of *ptsF* for cloning via *Xba*I |
| ptsF-Seq1 | GCAGGAAGCCACCACCGAG | Seq Primer ptsF Konstrukt |
| ptsF-Seq2 | GTTCAGGCTTCCTGTTGTACTTC | Seq Primer ptsF Konstrukt |
| lysC_fw | GAGGGATCCGAAAGGAGGCCCTTCAGGTGGCCCTGGTCGTA | Amplification of *lysC*$^{fbr}$ for cloning via *BamH*I / *Sac*I |
| lysC_rv | GAGGAGCTCTTAGCGTCCGGTGCCTG | Amplification of *lysC*$^{fbr}$ for cloning via *BamH*I / *Sac*I |
| FruK1_A | GTGACAACCGAAACAGTGCG | Amplification of *fruK1* upstream region for deletion |
| FruK1_B | CCCATCCACTAAACTTAAACAGGTGAATGTGATGATCATGGGGTTAC | Amplification of *fruK1* upstream region for deletion |
| FruK1_C | TGTTTAAGTTTAGTGGATGGGGTCACCCAAGTCAAAGGATTGAAAG | Amplification of *fruK1* downstream region for deletion |
| FruK1_D | CTTCGGATCGACTGGGGTG | Amplification of *fruK1* downstream region for deletion |
| FruK1_ver_f | TGGTCAAAAACCAGTTTCCCG | Verification of *fruK1* deletion |
| FruK1_ ver_r | ACTGGCTTGCCAGCGAAAC | Verification of *fruK1* deletion |
| FruK1_seq_fw | GTCGACGATTTCCTGCTCGG | Sequencing of deletion construct for *fruK1* deletion |
| FruK1_seq_rev | GGTGGACTGTTGTTACAAGTTCCC | Sequencing of deletion construct for *fruK1* deletion |
| FruK2_A | ATTCGATGTCACTGCAGAGACG | Amplification of *fruK2* upstream region for deletion |
| FruK2_B | CCCATCCACTAAACTTAAACAAGTGACTGTAAGAATCATTCTGCAA | Amplification of *fruK2* upstream region for deletion |
| FruK2_C | TGTTTAAGTTTAGTGGATGGGCTTCGGGCGGAGCACGTG | Amplification of *fruK2* downstream region for deletion |
| FruK2_D | ATGGCTGATGAGATGAACCCAG | Amplification of *fruK2* downstream region for deletion |
| FruK2_ver_f | CTTTTTGCTTTAAGGAGTGACATGTACG | Verification of *fruK2* deletion |
| FruK2_ver_r | GGCTGCGGTCTACTCAAAGG | Verification of *fruK2* deletion |
| Pfk_A | CGGGATCCCCAATGGAATGGTGCCCAGTGGGCGAA | Amplification of *pfk* upstream region for deletion |
| Pfk_B | CCCATCCACTAAACTTAAACAAATTCGCATGTCTTCCATATTAAACCCATCACAACACCCGC | Amplification of *pfk* upstream region for deletion |
| Pfk_C | TGTTTAAGTTTAGTGGATGGGGAACGCTGGGTTACTGCCCAGGCAATGTTT | Amplification of *pfk* downstream region for deletion |

| | | |
|---|---|---|
| Pfk_D | CGGGATCCCTCGCGTTGTTGGCCAATGCCCGG | Amplification of *pfk* downstream region for deletion |
| Pfk_ver_f | CAGTAGCTACTGCAGGACCCTTCTTTTC | Verification of *pfk* deletion |
| Pfk_ ver_r | CAACCACTCGGATGCGTGTGGATTC | Verification of *pfk* deletion |
| Hpr_ver_f | GTGCAGTCACTGATGCCTG | Verification of *hpr* deletion |
| Hpr_ver_r | GACATGAAAACCATGCACAGC | Verification of *hpr* deletion |
